# YeiE regulates YeiH to implement sulfite stress resistance in *Salmonella enterica* serotype Typhimurium

**DOI:** 10.1101/2025.07.27.662810

**Authors:** Trina L. Westerman, Elly Scheuerell, Vivian H. Nguyen, Kaiya Johnson, Michael McClelland, Johanna R. Elfenbein

## Abstract

Sulfites are toxic reactive sulfur species that are ubiquitous in nature. Bacteria are exposed to sulfites during the metabolism of sulfur-containing amino acids, during respiration using sulfur-containing electron acceptors, in host environments, and as food preservatives. Despite their broad distribution, the mechanisms bacteria use to resist the toxic effects of sulfites are poorly understood. Recent work showed that the LysR-type transcriptional regulator YeiE binds to sulfite to activate sulfite reduction gene expression. Using the model enteric pathogen, *Salmonella enterica* serotype Typhimurium, we show that YeiE is an autoregulator whose expression is enhanced by sulfite. We demonstrate here that the mechanism of sulfite stress resistance is distinct from pathways of sulfur assimilation for cysteine biogenesis. YeiE is necessary for survival in sulfite stress due to its regulation of the poorly characterized adjacent gene, *yeiH*. A *yeiH* deletion mutant is more sensitive to the growth-inhibiting effects of sulfite than a *yeiE* deletion mutant, demonstrating YeiH is the primary driver of sulfite stress resistance *in vitro*. This work provides a fundamental advance in understanding the sulfite stress response in bacteria with broad implications for food safety and host-pathogen interactions.

## Introduction

Sulfites are widely distributed reactive sulfur species that are toxic to all life forms. Sulfites are naturally present in many food products and have also been used for decades as food preservatives to reduce microbial growth and browning [1, 2]. All life forms produce sulfites endogenously during physiologic and pathologic states. Sulfites are byproducts of sulfated amino acid metabolism and in cysteine and methionine synthesis by many microbes [3–6]. Sulfites are also produced as a component of the innate immune response. Plasma sulfite levels are higher in people with bacterial pneumonia, and neutrophils produce sulfites in response to lipopolysaccharide, both *ex vivo* and in animal models of sepsis [7–10]. Furthermore, sulfites enhance neutrophil respiratory burst, suggesting sulfites may serve as a signaling molecule in the innate immune response against bacterial pathogens [11]. Given the prevalence of sulfites in the environment and within a host, bacterial pathogens must sense and respond to these compounds for survival.

Non-typhoidal *Salmonella enterica* (NTS) serotypes are a leading cause of bacterial foodborne gastroenteritis [12]. NTS are transmitted by ingesting contaminated food or water, or through direct contact with animals or people shedding the organism in feces. NTS invasion of the intestinal epithelium triggers neutrophil influx, causing severe neutrophilic inflammation and subsequent inflammatory diarrhea [13]. NTS may cross the gut barrier and replicate within professional phagocytes [14], causing life-threatening sepsis in immunosuppressed individuals [15]. Although NTS predominantly replicate in intracellular niches during systemic infection, neutrophils contribute to pathogen clearance in systemic tissues [16–18]. Considering the role of neutrophils in sulfite generation during infection, sulfite stress resistance is likely needed for *Salmonella* to withstand host antimicrobial defenses.

Bacteria have evolved efficient mechanisms to respond to environmental perturbations. LysR-type transcriptional regulators (LTTR) are the largest class of transcriptional regulatory proteins that respond to small-molecule signals. LTTRs possess an N-terminal winged helix-turn-helix DNA binding domain and a C-terminal effector binding domain [19]. YeiE is an LTTR linked to the pathogenesis of *Salmonella enterica*, *Cronobacter sakazakii,* and other pathogens [20–22]. The crystal structure of YeiE from *C. sakazakii* demonstrated that it binds to sulfite and controls expression of the sulfite reductase involved in sulfur assimilation for cysteine synthesis [23]. The *Salmonella enterica* serotype Typhimurium (*S*. Typhimurium) YeiE ortholog has 84% amino acid identity with *Cronobacter sakazakii*, leading us to hypothesize that YeiE regulates the sulfite stress response. Here, we demonstrate key regulatory features of YeiE in *S*. Typhimurium and show how it mediates survival in the face of sulfite stress.

## Results

### *yeiE* is needed for S. Typhimurium systemic infection

Prior work in our laboratory demonstrated that an *S*. Typhimurium Δ*yeiE* mutant was defective for swimming motility due to enhanced expression of a *Salmonella*-specific FlhD_4_C_2_ functional inhibitor (*STM1697*) [21, 24]. We observed reduced gut colonization in murine and bovine infection models and linked reduced gut colonization in the Δ*yeiE* mutant to altered flagella-mediated motility. Given that there are global regulatory targets of YeiE in *E. coli* [25], we hypothesized that YeiE would regulate expression of additional processes needed for *Salmonella* pathogenesis. To test this hypothesis, we used a murine sepsis model in which mice are inoculated with *Salmonella* intraperitoneally to avoid infection bottlenecks in the gut. Unlike in enteric infection, mutants lacking flagella-mediated motility are fully virulent in the murine sepsis model [26, 27]. Using competitive infections, we tested the impact of the Δ*yeiE* mutation compared to the isogenic WT in both male and female mice. We observed no differences in competitive index (CI) between sexes, allowing us to group data from both sexes. The Δ*yeiE* mutant had a competitive disadvantage compared with the WT organism in both the spleen and liver (Fig 1 and Table S1). These data suggest *yeiE* regulates processes in addition to flagella-mediated motility that are important for salmonellosis.

**Fig 1:**
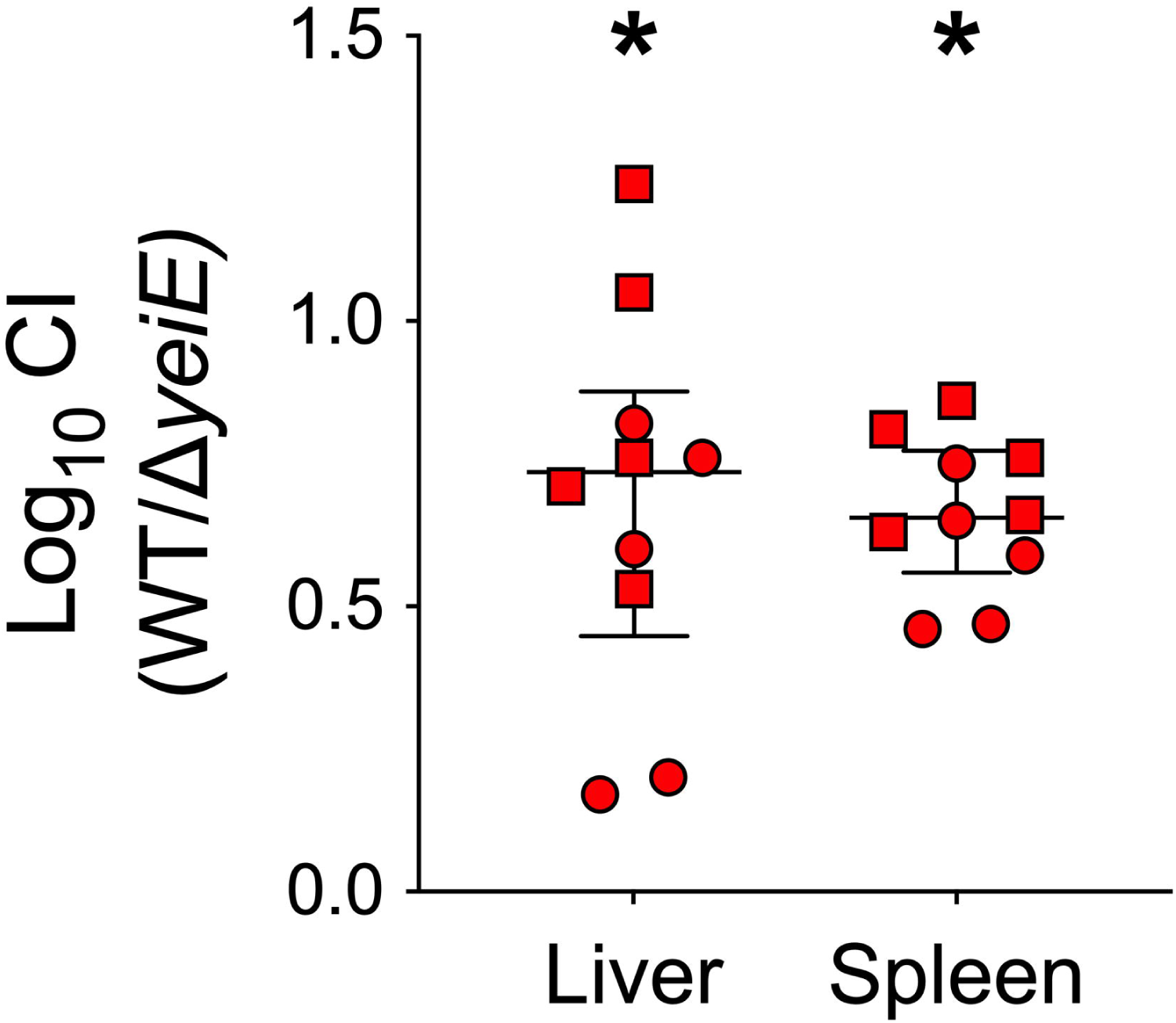
*yeiE* is necessary in systemic salmonellosis. Adult BALB/cJ mice were inoculated with ∼10^4^ CFU of an equal mixture of the WT and Δ*yeiE* mutant by intraperitoneal injection and euthanized 2 days post-infection. Competitive index (CI) is the ratio of WT to mutant after infection normalized to the inoculum ratio. Data from two independent experiments with female (circles) and male (squares) mice are displayed with median and interquartile range indicated. Statistical significance (*) was determined by Mann-Whitney test with Holm-Šídák’s correction for multiple comparisons.

### *yeiE* is required for survival in sulfite stress conditions

The crystal structure of YeiE from *C. sakazakii* showed that sulfite binds the YeiE effector binding domain, and a Δ*yeiE* mutant grew poorly in the presence of sulfite in both rich and minimal media [23]. We hypothesized that *yeiE* would also be needed for *S.* Typhimurium to grow in conditions of sulfite excess. We grew the WT and Δ*yeiE* mutant in rich media containing added sulfite. We found a severe growth defect for the Δ*yeiE* mutant grown in excess sulfite (5-40 mM; Fig 2A). The Δ*yeiE* mutant had a prolonged lag phase and truncated exponential phase when compared with the growth of the WT exposed to sulfite. The Δ*yeiE* mutant growth defect was sulfite-specific, as we did not observe any changes in growth when it was treated with similar sulfate concentrations (Fig 2B). We measured the area under the curve (AUC) in each treatment to quantify the impact of sulfite on total growth over the 12-hour experiment. Total growth of the WT was reduced when sulfite was added at ≥10 mM (Fig 2C). The Δ*yeiE* mutant grew poorly at 5mM sulfite with a further disadvantage at 10mM but no additional growth disadvantage at higher sulfite concentrations (Fig 2C). The complemented Δ*yeiE* mutant grew to the same density as the WT organism by the end of the experiment, though it displayed a biphasic growth pattern (Fig 2D). Total growth of the complemented Δ*yeiE* mutant was significantly improved when compared with the Δ*yeiE* mutant bearing an empty plasmid (Fig 2E), linking sulfite sensitivity phenotype to genotype.

**Fig 2:**
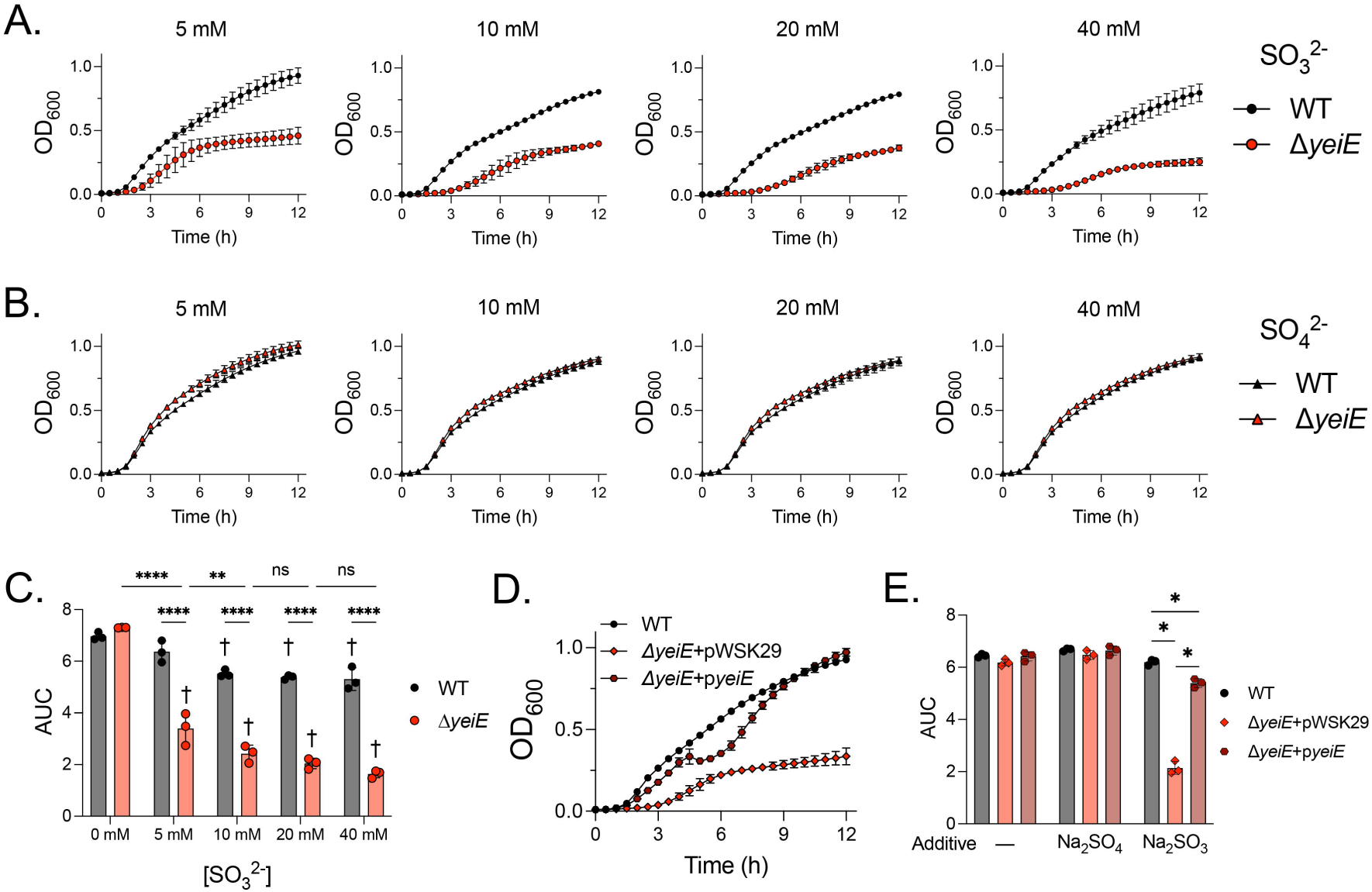
*yeiE* mediates survival in sulfite stress conditions. (A-B) The WT and Δ*yeiE* mutant were grown aerobically in Lennox broth with the indicated concentrations of (A) sulfite or (B) sulfate added. (C) The area under the curve (AUC) was determined from individual growth curves in sulfite (A). Statistical significance was determined by two-way ANOVA with Tukey’s multiple comparisons test. * indicates differences between the indicated strains or treatments and † indicates differences within a strain from 0 mM. ns – not significant. (D-E) The WT, Δ*yeiE* mutant with empty vector (Δ*yeiE*+pWSK29), and complemented Δ*yeiE* mutant (Δ*yeiE*+p*yeiE*) were grown aerobically in Lennox broth with (D) 5 mM sulfite, or (E) 5 mM sulfate, or untreated media. (E) The AUC from (D). Statistical significance was determined between strains by one-way ANOVA with Šidák’s multiple comparisons test. Data are mean +/- SD of blanked OD_600_ measurements from 3 independent experiments.

Since YeiE regulates the sulfite reductase encoded by *cysJI* in both *E. coli* and *C. sakazakii* [23, 25], we hypothesized that *yeiE* regulation of the assimilatory sulfite reductase (*cysJI*) would mediate survival in sulfite stress conditions. First, we tested the impact of *yeiE* on growth in minimal media because the *C. sakazakii* Δ*yeiE* mutant grew poorly in M9-sulfite [23]. Consistent with our prior work [21], the Δ*yeiE* mutant grew normally in M9-sulfate, indicating the pathway to produce cysteine from inorganic sulfur is intact in the Δ*yeiE* mutant (Fig 3A). However, the Δ*yeiE* mutant did not grow in M9-sulfite. As expected, the Δ*cysJ* mutant could not grow in minimal media with either sulfite or sulfate as sulfur source, indicating it is indeed a cysteine auxotroph. In contrast, the Δ*cysJ* mutant grew like the WT in sulfite stress in rich media (Fig 3B). Since the *S*. Typhimurium genome also encodes an anaerobic sulfite reductase (*asrABC*) that can transcomplement a sulfite reductase mutant in anaerobic conditions [28], we made a double Δ*cysJ*Δ*asrA* mutant and tested its growth in rich media with excess sulfite. Like the Δ*cysJ* mutant, the Δ*cysJ*Δ*asrA* mutant grew normally in high sulfite conditions in rich media (Fig 3B). The double mutant did not grow in M9-sulfite anaerobically. In contrast, the Δ*cysJ* single mutant grew, but not as efficiently as the WT in M9-sulfite, verifying that the anaerobic sulfite reductase can transcomplement the assimilatory sulfite reductase anaerobically in minimal media with sulfite as a sulfur source (Fig 3C). Because the Δ*cysJ*Δ*asrA* mutant does not phenocopy the Δ*yeiE* mutant in sulfite stress in rich media, these data suggest *yeiE* regulation of the sulfite reductases is not the primary mechanism mediating sulfite stress resistance in *S*. Typhimurium.

**Fig 3:**
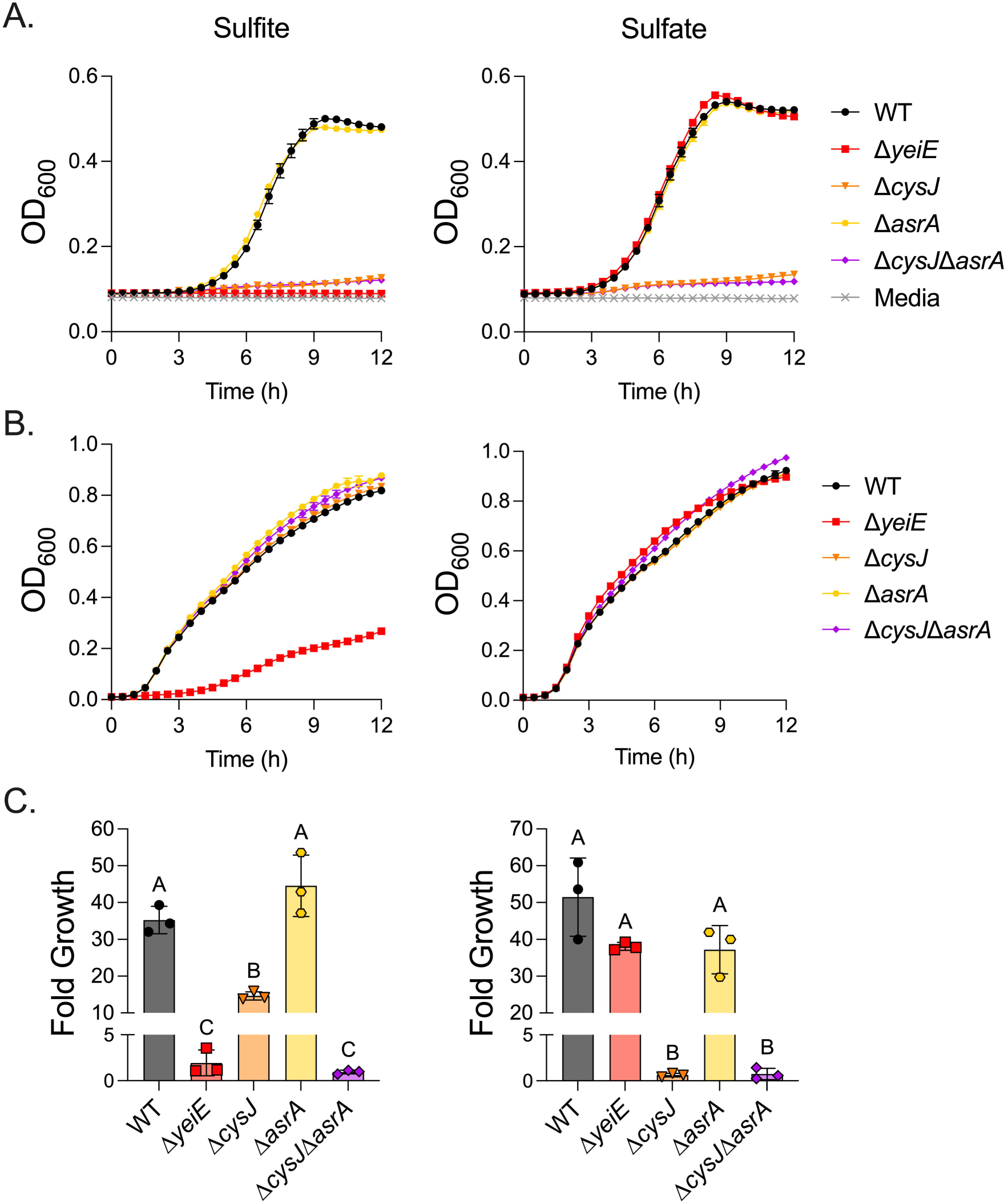
Sulfite reductases are dispensable for survival in sulfite stress conditions. (A) The WT, Δ*yeiE*, Δ*cysJ*, Δ*asrA*, and Δ*cysJ*Δ*asrA* mutants were grown aerobically in M9 with 1 mM sulfite or 1 mM sulfate as sulfur source. Uninoculated media is shown as the limit of detection of the assay, with raw measurements shown. (B) Bacteria were grown aerobically in Lennox broth with 10 mM sulfite or 10 mM sulfate. Blanked data are shown. (C) Bacteria were anaerobically grown for 24 hours in M9 with 1 mM sulfite or 1 mM sulfate as sulfur source. Fold growth is final CFU/mL normalized to input CFU/mL. Bars with different letters are significantly different by one-way ANOVA with Tukey’s correction for multiple comparisons. Mean +/- SD, n=3.

### YeiE is an autoregulator whose expression is enhanced by sulfite

In *E. coli*, YeiE has a strong ChIP binding peak to its own regulatory region [25]. To verify YeiE can bind to its own regulatory region in *S*. Typhimurium, we performed an EMSA using purified rYeiE and P*_yeiE_* DNA. We used P*_flhD_* DNA as a negative control, as *flhD* expression was not impacted by *yeiE* deletion in our prior study [21]. We observed a concentration-dependent gel shift for rYeiE+P*_yeiE_*, but not for P*_flhD_*, indicating YeiE binds to its own regulatory region in a dose-dependent and sequence-specific manner (Fig 4A). Next, we sought to determine how *yeiE* impacts its own expression using a transcriptional reporter with P*_yeiE_* driving luciferase expression. P*_yeiE_* activity was consistently detected in the WT background above control beginning at 1.5 hours (down arrow) and persisting for the remainder of the experiment (Fig 4B). In addition, P*_yeiE_* activity was always significantly higher in the Δ*yeiE* mutant background. These data demonstrate YeiE represses its own expression.

**Fig 4:**
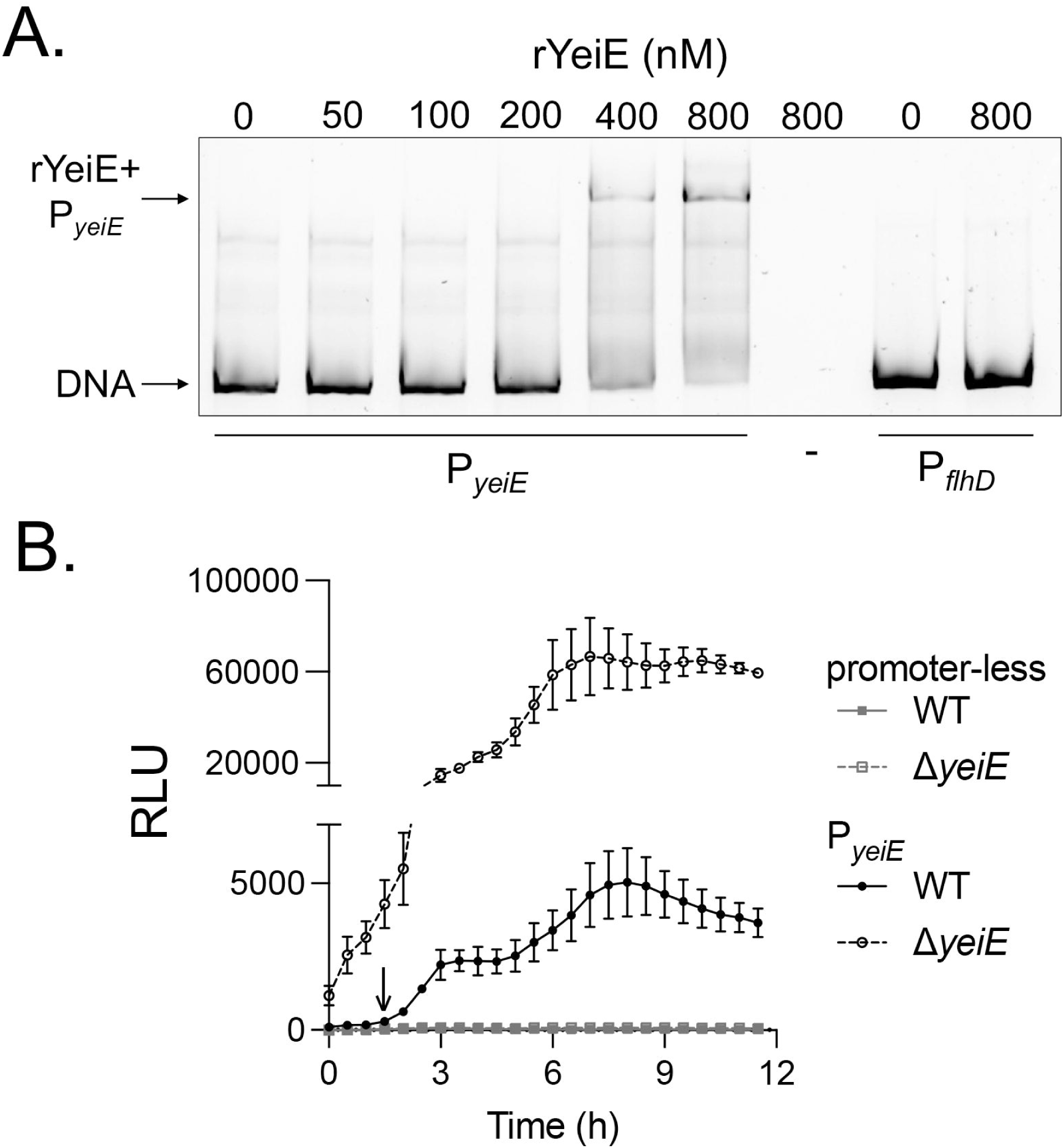
YeiE is an autorepressor. (A) EMSA of purified rYeiE (YeiE-strII) at the indicated concentrations was incubated with 40 ng DNA from the regulatory regions of the indicated genes. DNA-protein complexes were separated on a native 6% polyacrylamide gel with in-gel SYBR green staining. (B) The WT and Δ*yeiE* mutant, each bearing a plasmid with promoter-less *luxCDABE* (pCS26-Pac) or with the P*_yeiE_* transcriptional reporter (pCS26-Pac::P*_yeiE_-luxCDABE*) were grown aerobically in Lennox broth. Downward arrow is the time at which P*_yeiE_* RLU consistently increased above promoter-less plasmid in the WT by repeated-measures two-way ANOVA with Tukey’s correction for multiple comparisons. P*_yeiE_* expression was significantly higher in the Δ*yeiE* mutant than in the WT at all times tested. Blanked relative luminescence units (RLU) from 3 independent experiments (mean +/- SD) are shown.

Next, we tested the impact of sulfite on *yeiE* expression. We grew bacteria to early exponential phase growth (1.5 hours) to activate *yeiE* expression and then added sulfite or control compounds to the cells. We compared the relative luminescence units (RLU) to OD_600_ to determine P*_yeiE_* activity on a per-cell basis. Sulfite significantly increased P*_yeiE_* activity above all other groups from 1-1.5 hours post-treatment (Fig 5A). We hypothesized that *yeiE* would be responsible for the increased P*_yeiE_* activity upon sulfite treatment. Because P*_yeiE_* activity is higher in the Δ*yeiE* mutant (Fig 4B), we normalized P*_yeiE_* per-cell activity after one hour of sulfite treatment to its activity before treatment in the WT and Δ*yeiE* mutant backgrounds. Sulfite significantly increased P*_yeiE_* activity in the WT but had no impact on the Δ*yeiE* mutant (Fig 5B). These data demonstrate that it is the interaction of YeiE with sulfite that activates P*_yeiE_* activity. To verify whether changes in P*_yeiE_* activity translate to YeiE protein abundance, we determined the impact of sulfite on YeiE-3xFLAG abundance using a chromosomal translational reporter. We observed a more than 3-fold increase in YeiE-3xFLAG abundance within 1.5 hours after sulfite treatment, while YeiE-3xFLAG increased ∼1.6 fold in untreated and sulfate control conditions (Fig 5C-D). The abundance of rYeiE in sulfite-treated cells was significantly higher than in untreated cells within 15 minutes of treatment and significantly higher than in sulfate-treated cells by 45 minutes. Together, these data demonstrate that sulfite enhances YeiE expression through enhanced activity of P*_yeiE_*.

**Fig 5:**
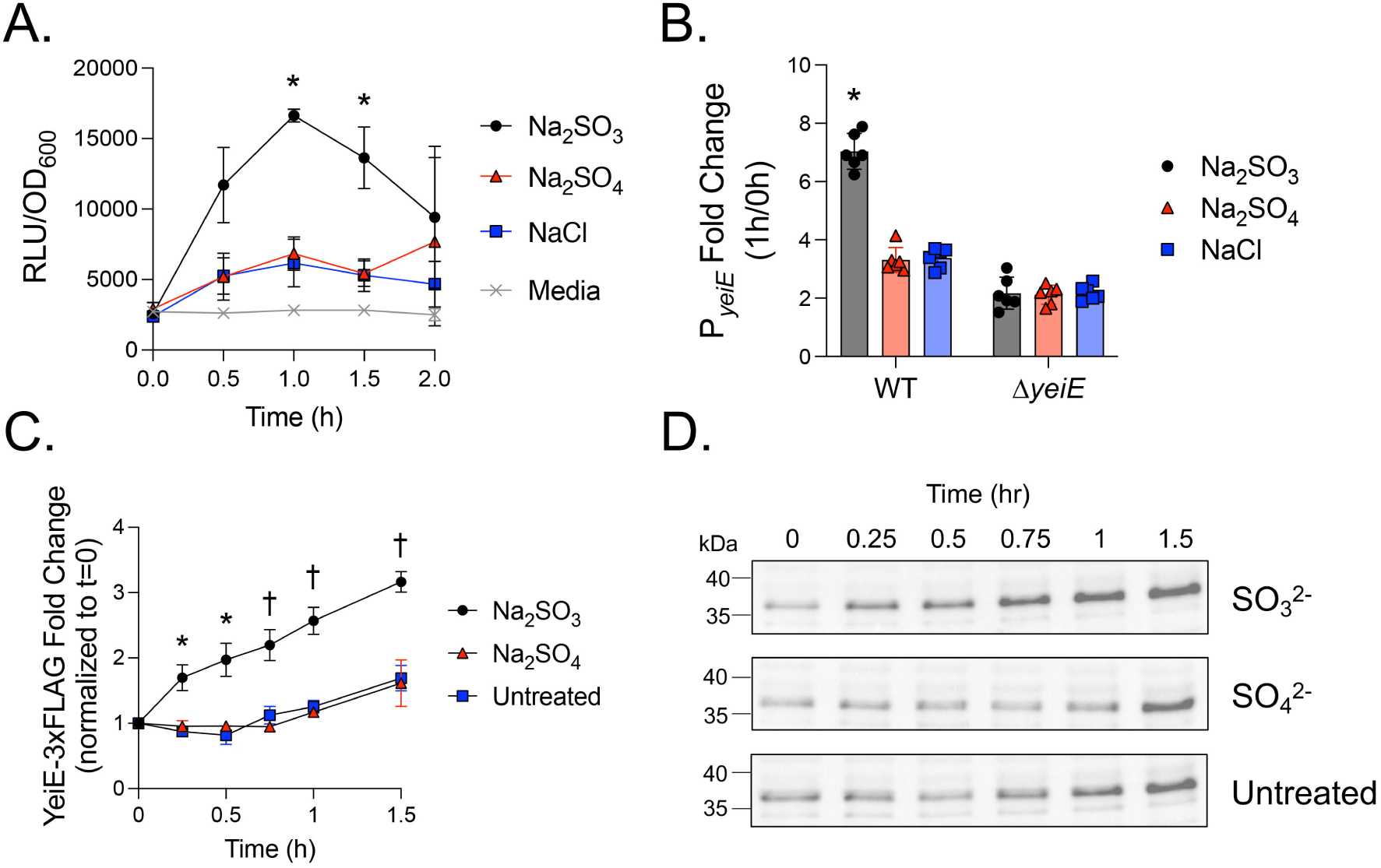
Sulfite enhances *yeiE* expression. (A-D) Bacteria grown aerobically to the early exponential phase in Lennox broth were treated with the indicated concentrations of sulfite, sulfate, sodium chloride, or not treated. (A) The WT bearing the P*_yeiE_* transcriptional reporter was treated with 1 mM sulfite or sulfate, or 2 mM sodium chloride. RLU and OD_600_ were determined at the indicated times post-treatment. Untreated media represents the lower limit of detection of the assay. * indicates significant difference between sulfite and both sulfate and sodium chloride treatments by repeated measures two-way ANOVA with Geisser-Greenhouse correction and Tukey’s multiple comparisons test. Mean +/- SD, n=3. (B) WT or Δ*yeiE* mutant, each bearing P*_yeiE_* transcriptional reporter, were treated with 2 mM sulfite, sulfate, or 4 mM sodium chloride. RLU/OD_600_ after 1 hour treatment was normalized to pre-treatment (t=0h) for each strain and treatment. * indicates significant differences between sulfite and both sulfate and sodium chloride treatments by two-way ANOVA with Tukey’s multiple comparisons test. Mean +/- SD, n=6. (C) Bacteria with YeiE-3xFLAG chromosomal fusion were treated with 1 mM sulfite or sulfate. Cultures were normalized by OD_600,_ and proteins were separated on 10% SDS-PAGE gels. YeiE-3xFLAG (∼36 kDa) intensity from western blot was normalized to total protein loading control. Fold change in normalized YeiE-3xFLAG intensity at each time was compared with pre-treatment (t=0h). * indicates significant difference from untreated and † indicates difference from both sulfate-treated and untreated cultures by repeated measures two-way ANOVA with Geisser-Greenhouse correction and Tukey’s multiple comparisons test. Mean +/- SD, n=3. (D) Representative western blot of YeiE-3xFLAG from (C).

*yeiH* is essential for sulfite stress resistance.

*yeiH* is neighboring and divergently expressed from *yeiE,* with 120 bp separating their coding sequence start codons. We showed rYeiE binds to P*_yeiE_* (Fig 4A), which included the *yeiH* regulatory region and 120 bp into the *yeiH* coding sequence (Fig S2). Therefore, we hypothesized *yeiH* would be regulated by YeiE. First, we tested the impact of sulfite on *yeiH* expression using a transcriptional reporter plasmid with P*_yeiH_* driving luciferase expression. Sulfite treatment in the early exponential growth phase increased per-cell P*_yeiH_* activity within 30 minutes in a dose-dependent manner (Fig 6A). This effect was specific to sulfite, as P*_yeiH_* activity in untreated or sulfate-treated cells was always below the limit of detection (uninoculated media). We treated the WT and Δ*yeiE* mutant, each bearing the P*_yeiH_* transcriptional reporter, with sulfite to establish whether sulfite-mediated P*_yeiH_* activation requires *yeiE*. At 30 minutes after sulfite treatment, we observed a significant increase in P*_yeiH_* activity in the WT but no change in expression in the Δ*yeiE* mutant background (Fig 6B). These data demonstrate that YeiE sensing sulfite is essential to activate *yeiH* transcription.

**Fig 6.**
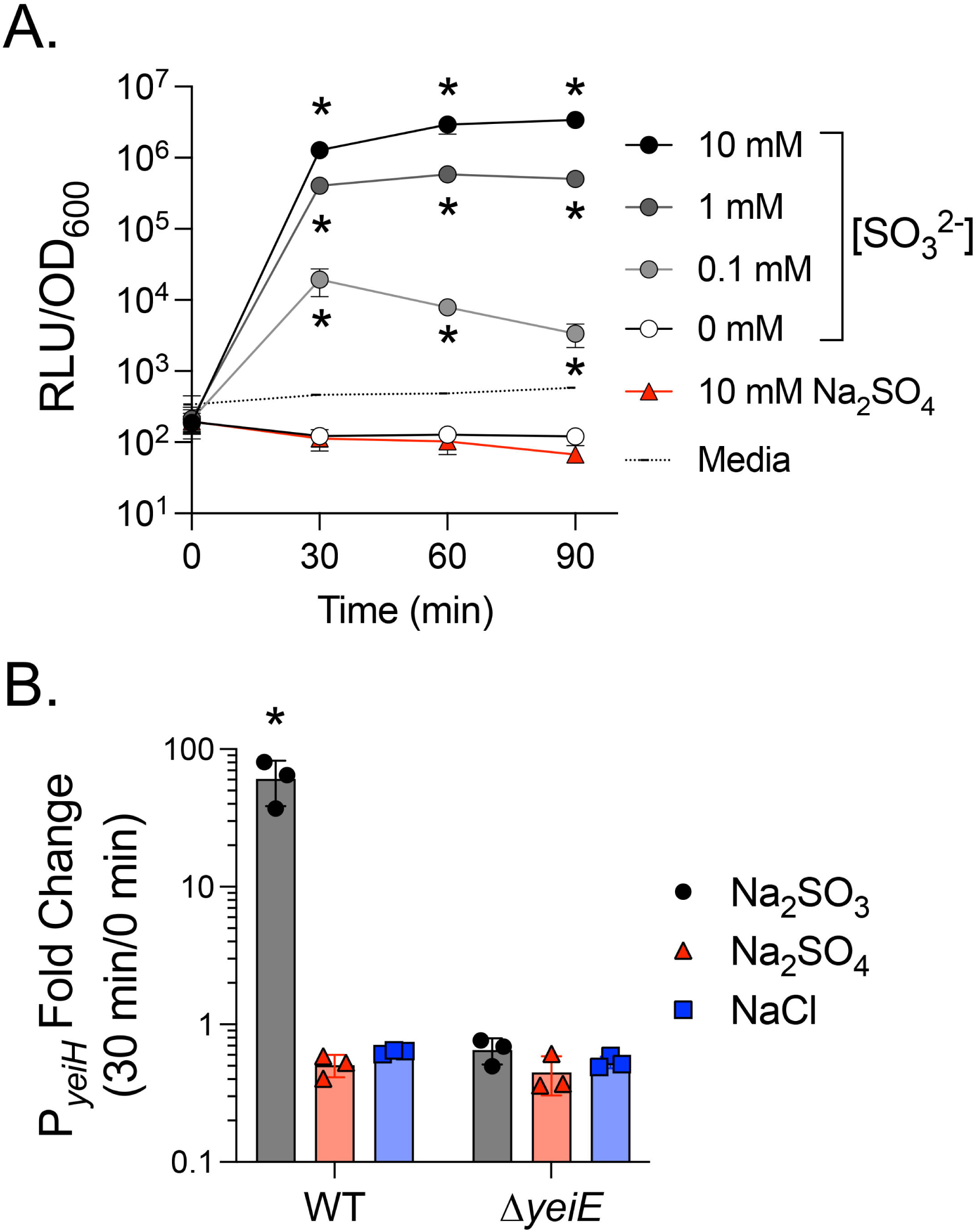
P*_yeiH_* expression is activated by YeiE in response to sulfite. (A) The WT bearing P*_yeiH_* transcriptional reporter, grown aerobically to early exponential phase in Lennox broth, was treated with the indicated concentrations of sulfite, sulfate, or not treated. RLU and OD_600_ were determined at the indicated times post-treatment. Untreated media represents the lower limit of detection of the assay. * indicates significant difference between the indicated group and uninoculated media by repeated measures two-way ANOVA with Geisser-Greenhouse correction and Tukey’s multiple comparisons test. Mean +/- SD, n=4. (B) The WT and Δ*yeiE* mutant, each bearing P*_yeiH_* transcriptional reporter, grown as in (A), were treated with 0.1 mM sulfite or sulfate or 0.2 mM sodium chloride. RLU/OD_600_ after 30 minutes of treatment was normalized to pre-treatment (t=0h) for each strain and treatment. * indicates significant differences between sulfite and both sulfate and sodium chloride treatments by two-way ANOVA with Tukey’s multiple comparisons test. Mean +/- SD, n=3.

To test our hypothesis that *yeiH* would be needed for *S*. Typhimurium to survive sulfite stress, we compared the growth of the WT and Δ*yeiE* and Δ*yeiH* mutants in rich media supplemented with sulfite. The Δ*yeiH* mutant had a severe growth defect in sulfite, barely replicating within the 12-hour experiment (Fig 7A). The growth defect was sulfite-specific, as the Δ*yeiH* mutant grew normally in untreated media and in sulfate-treated media. We also noted the Δ*yeiH* mutant was more attenuated for growth than the Δ*yeiE* mutant when treated with sulfite. We tested a range of sulfite concentrations to determine the relative sensitivity of the Δ*yeiE* and Δ*yeiH* mutants to sulfite (Fig 7B). The Δ*yeiH* mutant was 4 times more sulfite sensitive than the Δ*yeiE* mutant (0.5 mM vs 2 mM) and 20 times more sulfite sensitive than the WT (0.5 mM vs 10 mM (Fig 2C)). The complemented Δ*yeiH* mutant (Fig 7C) grew like the WT in sulfite-supplemented media, definitively linking *yeiH* with the sulfite sensitivity phenotype.

**Fig 7.**
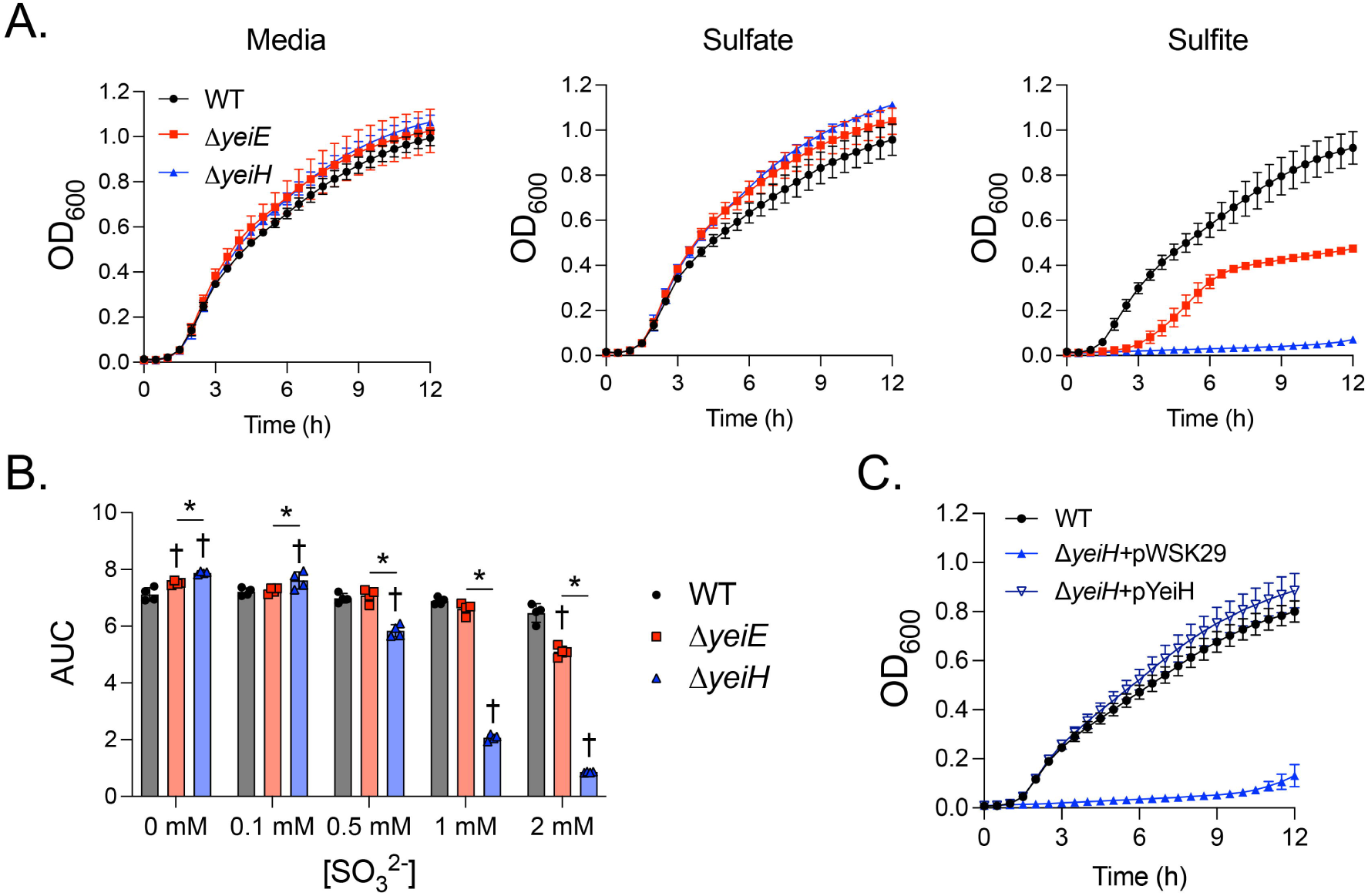
*yeiH* is key to *S*. Typhimurium sulfite stress resistance. (A) Bacteria were grown aerobically in Lennox broth with 5 mM sulfite, sulfate, or untreated. Mean +/- SD of blanked OD_600_ data, n=3. (B) AUC for each biological replicate of bacteria treated with the indicated concentrations of sulfite. * indicates significant differences between the Δ*yeiE* and Δ*yeiH* mutants within a treatment, and † indicates significant differences between the indicated strain and the WT within a treatment by repeated measures two-way ANOVA with Tukey’s multiple comparisons test. Mean +/- SD, n=4. (C) The WT, Δ*yeiH* mutant with empty plasmid (pWSK29) and complemented Δ*yeiH* mutant (p*yeiH*) were grown as in (A). Mean +/- SD of blanked OD_600_ data, n=3.

Next, we investigated whether the lack of increase in cell density of the sulfite-treated Δ*yeiH* mutant was due to death or inhibition of growth. We grew bacteria to the early exponential phase and then treated them with 10 mM sulfite, the lowest dose that attenuated the growth of the WT bacterium, or with 10 mM sulfate. Samples were taken at regular intervals, diluted, and plated to determine viable CFU. Sulfite treatment had no impact on the growth of the WT but stopped the growth of both the Δ*yeiE* and Δ*yeiH* mutants (Fig 8A). The bacteriostatic effect of sulfite persisted for 24 hours for the Δ*yeiH* mutant, but the Δ*yeiE* mutant was able to achieve a similar cell density as the WT within 24 hours. Although we observed no growth defect for the sulfite reductase mutant (Δ*cysJ*Δ*asrA*) in sulfite-treated rich media, we hypothesized that the sulfite reductases may be activated in a *yeiE*-independent manner to enhance late growth of the Δ*yeiE* mutant. When a triple Δ*yeiE*Δ*cysJ*Δ*asrA* mutant was treated with sulfite, its growth was identical to that of the Δ*yeiE* mutant, indicating the late growth recovery was independent of sulfite reduction (Fig 8B). Though sulfite reduction did not impact the growth recovery of the Δ*yeiE* mutant, we hypothesized that removal of sulfite reduction would render sulfite bactericidal for the Δ*yeiH* mutant. However, the triple Δ*yeiH*Δ*cysJ*Δ*asrA* mutant had no further alteration to viability upon sulfite treatment when compared with the Δ*yeiH* mutant (Fig 8C). When taken together, these data demonstrate that *yeiH* is the primary driver necessary to resist the bacteriostatic impacts of sulfite.

**Fig 8.**
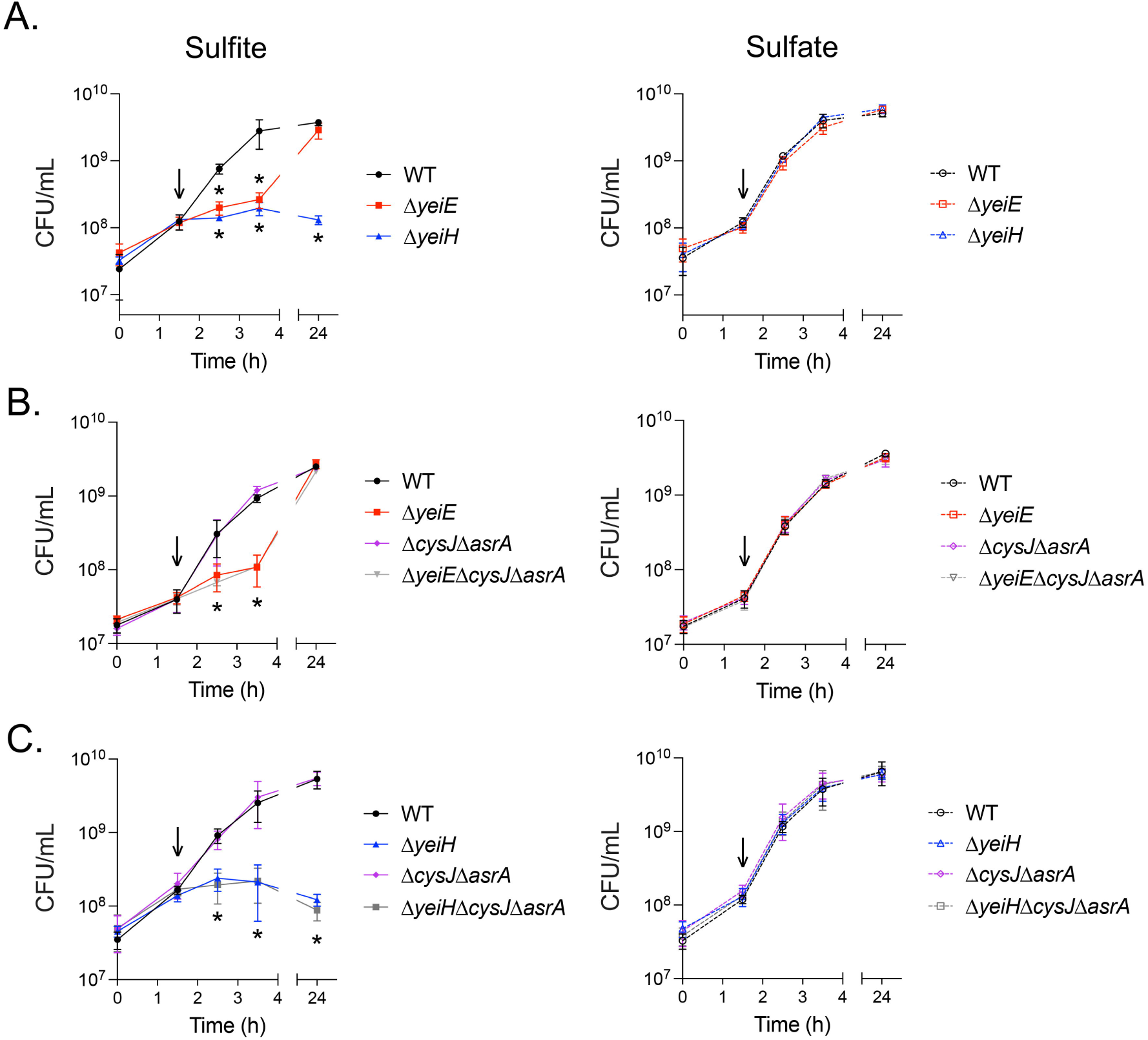
Sulfite reduction does not influence the viability of *yeiE* or *yeiH* mutants during sulfite stress. The indicated bacterial strains were grown aerobically in Lennox broth to early exponential phase at which time 10 mM sulfite or sulfate was added (downward arrow). Aliquots of bacteria were removed at the indicated times, diluted, and plated to establish CFU/mL. * indicates significant difference from the WT treated with sulfite at the indicated time by repeated measures two-way ANOVA with Dunnet’s correction for multiple comparisons. Mean +/- SD, n=3.

## Discussion

Structural and thermodynamic studies of YeiE and its homologs demonstrated that it binds sulfite [23, 29]. This information led to the conclusion that YeiE homologs regulate the expression of sulfur homeostasis. Here, we add functional and phenotypic data to show the role of YeiE in controlling the response to sulfite stress. We demonstrate that YeiE binds to its own promoter DNA to repress its own transcription in the absence of sulfite. Autorepression in the absence of ligand is postulated to allow LTTRs to respond to external stimuli quickly [19]. Consistent with this observation, we show that YeiE rapidly activates expression from its own promoter and the neighboring *yeiH* promoter when sulfite is present. Enhanced transcription leads to increased YeiE abundance, indicating YeiE accumulates in response to sulfite stress. In turn, YeiE responds to sulfite by activating *yeiH* transcription in a sulfite dose-dependent manner. During the completion of our work, it was shown that the *Pseudomonas aeruginosa* YeiE homolog FinR, binds to its own promoter and sulfite influences the specific DNA sequence to which the protein binds [29]. Our data in S. Typhimurium are consistent with the model that YeiE binding to specific DNA sequences in the *yeiE-yeiH* intergenic region changes upon sulfite binding to activate transcription from both divergent promoters. Future work is needed to establish how sulfite binding influences expression of genes under transcriptional control of YeiE.

Our work shows that *yeiH* is the key to sulfite stress resistance. Though *yeiH* is poorly expressed during aerobic growth in Lennox broth [30], *yeiH* promoter activity increases in a YeiE-dependent and sulfite dose-dependent manner, consistent with a specific role for *yeiH* in sulfite stress. The Δ*yeiH* mutant is highly sensitive to sulfite, with a growth defect at 4x lower sulfite concentration than the Δ*yeiE* mutant. In addition, the Δ*yeiE* mutant completely recovers to WT levels by 24 hours of aerobic growth in sulfite stress, whereas the Δ*yeiH* mutant fails to recover. Removing both annotated sulfite reductases (Δ*cysJΔasrA*) from the Δ*yeiE* mutant did not influence its growth recovery, providing evidence that sulfite reduction does not contribute to growth in aerobic sulfite stress *in vitro*. We hypothesize that *yeiH* activation in a *yeiE*-independent manner during late growth in sulfite stress allows the Δ*yeiE* mutant to recover but not the Δ*yeiH* mutant.

*yeiH* is highly conserved among eubacteria with some orthologs annotated as putative sulfate exporters, although it remains a poorly characterized gene [31]. YeiH is predicted to be a membrane protein with nine transmembrane helices, a pair of broken helices that enter and exit on the same side of the membrane, and a periplasmic C-terminus [32–34]. YeiH has structural similarity to a Na^+^/H^+^ antiporter and a sodium-dependent glutamate transporter [32, 33], suggesting it has a transport function. Aside from protein structural predictions, very little is known about *yeiH* function. An *E. coli* MG1655 Δ*yeiH* mutant grows poorly in the presence of glutathione [35]. These data, together with our data showing the essential role of *yeiH* in sulfite stress resistance, suggest an important role for *yeiH* in sulfur homeostasis. Future work will elucidate the mechanism by which *yeiH* mediates sulfite stress resistance in S. Typhimurium and other enteric pathogens.

We used growth in both minimal and rich media with sulfite to disentangle the potentially overlapping role of *yeiE* in sulfite stress with that of sulfur metabolism. YeiE regulates the *cysJI* sulfite reductase in both *E. coli* and *C. sakazakii* [23, 25]. Using minimal media, we verified published work demonstrating that the *S*. Typhimurium Δ*yeiE* mutant has normal growth characteristics in minimal media with sulfate [21]. These data indicate that *yeiE* does not contribute to cysteine biosynthesis in sulfur-limiting conditions in *S*. Typhimurium. In *E. coli* and *S*. Typhimurium, the LTTR CysB responds to N-acetylserine levels to activate expression of genes for assimilatory sulfate metabolism (reviewed in [36]). However, the *C. sakazakii* Δ*yeiE* mutant has a slight growth disadvantage in minimal media with sulfate as the sulfur source [23], suggesting *yeiE* may contribute to cysteine biosynthesis in *C. sakazakii*. *C. sakazakii* has a *cysB* homolog [37], but it is possible that the regulation of assimilatory sulfate metabolism differs between the two organisms. It is also possible that the amount of sulfite generated during sulfate assimilation overwhelms the *C. sakazakii ΔyeiE* mutant defenses. Despite the differences in growth characteristics with sulfate, both organisms required *yeiE* and the *cysJI* sulfite reductase to grow aerobically in minimal media with added sulfite, suggesting a conserved mechanism for sulfite metabolism for aerobic cysteine biosynthesis.

*S*. Typhimurium also encodes an anaerobic sulfite reductase (*asrABC*) that can transcomplement a sulfite reductase mutant in anaerobic conditions [28, 38]. Therefore, we tested the sulfite sensitivity of a double Δ*cysJ*Δ*asrA* mutant, which we verified was unable to grow in minimal media with sulfite or sulfate aerobically. Anaerobically, the Δ*cysJ* mutant was capable of growth in minimal media with sulfite, but the Δ*cysJ*Δ*asrA* mutant was not, verifying anaerobic transcomplementation of the assimilatory sulfite reductase mutant. To our surprise, the Δ*cysJ* mutant also grew poorly in anaerobic minimal media with sulfate, indicating the anaerobic sulfite reductase was unable to transcomplement growth with sulfate. Prior work suggested a Δ*cysJ* deletion mutant had defective *cysH* expression due to polar effects, but the mutant was mapped to the promoter in addition to the *cysJ* coding region [39]. Though our Δ*cysJ* mutant has the native promoter and first 30 nt of the coding region intact [40], this is followed by a cassette containing a promoter and Kanamycin resistance gene in the sense strand, so we cannot rule out polarity on downstream gene expression. Furthermore, the minimal media we used lacked any transition metals, whereas previous work provided them [41], so it is possible media differences could account for the differences in anaerobic sulfate assimilation of the Δ*cysJ* mutant. Expression of the *asr* operon is induced by sulfite in the absence of oxygen [38], leading us to hypothesize that YeiE senses sulfite to activate anaerobic expression of the *asr* operon, but sulfate assimilation generates insufficient sulfite to activate *asr* expression. Further work is needed to elucidate the mechanism of regulation of the anaerobic sulfite reductase.

Pathogens may experience sulfite stress in several ways *in vivo*. There are a wide variety of inorganic and organic sulfur species in the gastrointestinal tract, both during homeostasis and in disease. Sulfites can be generated by host and microbial metabolism of inorganic and organic sulfur compounds [4]. Thus, enteric pathogens will encounter sulfites upon entry into a normal gut ecosystem. During intestinal inflammation, such as in enteric salmonellosis, the neutrophil oxidative burst oxidizes thiosulfate normally present in the gut to tetrathionate. *Salmonella* uses the tetrathionate reductase to reduce tetrathionate to thiosulfate for anaerobic respiration [42]. Sulfite is produced by the reduction of thiosulfate [43]. In addition, sulfites can be produced by reactions of inorganic sulfur compounds with reactive oxygen species produced by activated neutrophils in tissues [7, 9]. Our data show the *yeiE*-mediated sulfite stress response is needed during systemic infection, but whether the sulfite stress response is activated during tetrathionate-thiosulfate reduction in the gut is currently unknown. This work provides the foundation for future studies to understand the impact of sulfite stress on host-pathogen interactions.

## Methods

### Bacterial strains, plasmids, and reagents

Bacteria are derivatives of *Salmonella enterica* serotype Typhimurium ATCC14028s and are presented in Table S2. The Δ*yeiH* mutation was retrieved from the *Salmonella* Typhimurium Single Gene Deletion library and moved into a clean genetic background using bacteriophage P22-mediated transduction, with mutation verified by PCR using primers flanking the gene of interest [40, 44, 45]. Antibiotic resistance cassettes were removed using *flp* recombinase [46]. All bacteria were stored at −80°C in 30% glycerol until use. Unless otherwise specified, bacteria were routinely grown at 37°C in Luria-Bertani broth (LB-Miller, BD Difco™) with agitation (225 rpm) or on LB agar (BD Difco™). The antibiotics nalidixic acid (50 mg/L), kanamycin (50 mg/L), carbenicillin (100 mg/L), and chloramphenicol (20 mg/L) were used when indicated.

Sodium sulfite (Na_2_SO_3_; VWR), sodium sulfate (Na_2_SO_4_; Thermo Scientific), and sodium chloride (NaCl; Fisher Scientific) were used at the concentrations indicated. Sodium sulfite and sodium sulfate were dissolved in deionized water to 1M and sodium chloride to 2M and filter-sterilized prior to dilution to final experimental concentrations. Stock solutions were stored at room temperature and used within 3 weeks of preparation. Synthetic nucleic acids were purchased from Integrated DNA Technologies.

### Plasmid construction

*yeiE* encoding a C-terminal Strep-tag® II with S-A linker (SAWSHPQFEK) was cloned into pBAD18-Cm [47] to create an inducible YeiE-Strep-tag® II for protein purification. Colony PCR was performed using Q5 DNA Polymerase (New England Biolabs) and primers 5yeiE-strKpnIpBAD18 and 3yeiE-strHindIIIpBAD18. Primer sequences and reaction information are available in Table S3. Following verification of product size and purity, the PCR product was purified (QIAquick® PCR Purification Kit, Qiagen), digested with KpnI and HindIII (New England Biolabs), and purified again. The pBAD18-Cm vector was sequentially digested with KpnI and HindIII and treated with recombinant Shrimp Alkaline Phosphatase (New England Biolabs). The insert was ligated into the cut vector using T4 DNA Ligase (New England Biolabs). The reaction mixture was transformed into E. coli DH5α by heat-shock and plated onto LB agar containing chloramphenicol. Colonies were purified twice. Plasmid was isolated (QIAprep® MiniPrep Kit, Qiagen) and insert verified by Sanger sequencing (Functional Biosciences). The pBAD18-Cm::*yeiE-strII* construct was transformed into *E. coli* strain BL21 (New England Biolabs) by heat shock.

Transcriptional reporter plasmids were generated from pCS26-Pac encoding a reporter-less *luxCDABE* [48]. Colony PCR using Q5 DNA Polymerase was used to amplify the *yeiE* regulatory region (P*_yeiE_*) of −518 to +50nt from the start codon using primers PyeiEFWD2XhoI and PyeiEREV2BamHI or the *yeiH* regulatory region (P*_yeiH_*) of −600 to +60nt from the start codon using primers PyeiHXhoIpCS26FWD and PyeiHBamHIpCS26REV (Table S3). The sequences of P*_yeiE_* and P*_yeiH_* are available in Fig S1. Product size and purity were established by agarose gel electrophoresis, and products were digested with XhoI and BamHI (New England Biolabs) followed by PCR purification. T4 DNA Ligase was used to ligate digested PCR products into pCS26-Pac digested with XhoI and BamHI and treated with recombinant Shrimp Alkaline Phosphatase. Ligation reactions were transformed into *E. coli* DH5α by heat shock and plated onto LB agar with kanamycin. Transformants were purified twice, and plasmids isolated. Plasmid sequences were verified by whole-plasmid sequencing (Plasmidsaurus). Sequence-verified plasmids were transformed into S. Typhimurium by electroporation. Transformants were purified twice and stored at −80°C until use. The stable plasmid pWSK29 was used to complement the Δ*yeiH* mutation *in trans* [49]. Colony PCR using Q5 DNA Polymerase was used to amplify *yeiH* with its native promoter using primers yeiHBamHIFWD and yeiHHindIIIREV (Table S3). Product size and purity were determined by agarose gel electrophoresis, and the product was digested with BamHI and HindIII. The product was ligated into pWSK29 sequentially digested with HindIII and BamHI and treated with recombinant Shrimp Alkaline Phosphatase using T4 DNA Ligase. Ligation reactions were transformed into *E. coli* DH5α by heat shock and plated onto LB with carbenicillin and X-gal (5- Bromo-4-Chloro-3-Indolyl β-D-Galactopyranoside; 40 µg/mL). Transformants were purified twice, plasmids isolated, and sequence verified by whole-plasmid sequencing. Plasmids were transformed into restriction minus, modification positive *S*. Typhimurium LB5000 by electroporation [50]. Colonies were purified twice, plasmids isolated, and transformed into the Δ*yeiH* mutant by electroporation. The complemented mutant was purified twice and stored at - 80°C until use.

### Bacterial growth curves

To measure the impact of sulfite on bacterial growth in rich media by optical density (OD_600_), overnight cultures were diluted 1:100 into Lennox broth (BD Difco™) containing the indicated concentrations of sodium sulfite, sodium sulfate, or no additive. Two hundred microliters of bacterial suspension was added to duplicate or triplicate wells of a clear, U-bottom 96-well plate. Uninoculated media with additives was added to plates to serve as media blanks. Plates were incubated in a plate reader (Tecan Infinite 200 PRO® MPlex) at 37°C with orbital shaking (3mm amplitude) and OD_600_ measured every 15 minutes. Data from duplicate or triplicate wells were averaged, and the average absorbance of the uninoculated media was subtracted from the absorbance reading at each data collection time to display blanked data. Every other data point of blanked OD_600_ data is shown for clarity. Each experiment was performed on at least three independent occasions.

To determine the impact of sulfite on growth in minimal media, overnight cultures were pelleted and washed twice in sulfur-free (SF) M9 minimal media (48 mM Na_2_HPO_4_, 22 mM KH_2_PO_4_, 9 mM NaCl, 19 mM NH_4_Cl, 0.1 mM CaCl_2_ and 1 mM MgCl_2_) to eliminate any remaining sulfur. For aerobic growth, bacteria were resuspended in SFM9 and diluted 1:100 into M9 with 1 mM Na_2_SO_4_ or Na_2_SO_3_ as the sulfur source. Bacterial suspensions were added to 96-well plates and grown as above. Data from duplicate wells were averaged, and every other data point is displayed for clarity. For anaerobic growth, washed bacteria were pelleted again and supernatants removed. Cell pellets were transferred into an anaerobic chamber with an atmosphere composed of 5% H_2_, 5% CO_2_, and 90% N_2_ (Shel Lab; Bactron EZ). Pellets were resuspended in SFM9 that was pre-reduced in the anaerobic chamber for a minimum of 12 hours. Cultures were then diluted 1:100 into 2 mL pre-reduced M9 supplemented with 1 mM Na_2_SO_4_ or Na_2_SO_3_ as the sulfur source. Cultures were grown standing at 37°C in the anaerobic chamber with 200 µL aliquots taken at 0 and 24 hours, serially diluted, and plated to determine colony-forming units (CFU)/mL. Each experiment was performed on three independent occasions.

To measure the impact of sulfite treatment on exponentially growing cells, overnight cultures were diluted 1:100 into 50 mL Lennox broth in a 250 mL flask. Bacteria were incubated at 37°C with agitation for 1.5 hours, at which time cultures were treated with sulfite or sulfate at the indicated concentrations. One milliliter of culture was removed at the indicated times, serially diluted, and plated to determine CFU/mL. Each experiment was performed on at least three independent occasions.

### Promoter activity measurements

Overnight bacterial cultures bearing the P*_yeiE_* transcriptional reporter plasmid or the promoter-less plasmid were diluted 1:100 into Lennox broth, pH 7.0, with added kanamycin. Two hundred microliters of bacteria were added to a sterile white, flat-bottomed 96-well plate in triplicate. Uninoculated media was added to triplicate wells as a blank for background luminescence. Plates were incubated in a plate reader (Tecan Infinite 200 PRO® MPlex) at 37°C with orbital shaking (4mm amplitude) and relative luminescence units (RLU) measured prior to incubation and every 15 minutes (1000 ms integration time). Data from triplicate wells was averaged, and the average RLU from uninoculated media was subtracted to generate blanked RLU measurements.

To measure the impact of sulfite on *yeiE* or *yeiH* promoter activity, overnight cultures from bacteria harboring P*_yeiE_* or P*_yeiH_* reporter plasmids were diluted 1:100 into 50 mL Lennox broth with kanamycin in 250 mL flasks. Bacteria were grown at 37°C with agitation for 1.5 hours, at which time sulfite or sulfate was added. At the indicated times, 1 mL of bacteria was removed for measurement of both OD_600_ and RLU (Tecan Infinite 200 PRO® MPlex) in duplicate measurements. Duplicate measurements were averaged, and the RLU was divided by the OD_600_ to estimate promoter activity on a per-cell basis. The RLU/OD_600_ of uninoculated media is presented as the lower limit of detection of the assay. Each experiment was performed on at least three independent occasions.

### Electrophoretic mobility shift assays

Recombinant YeiE-strII (rYeiE) was purified from *E. coli* BL21. Overnight cultures of BL21+pBAD18Cm::*yeiE-strII* were diluted 1:100 into LB containing chloramphenicol and 2% glucose. Bacteria were grown aerobically at 37°C until reaching an OD_600_ of 0.4-0.5. Cells were centrifuged, supernatant removed, resuspended in LB with chloramphenicol and 0.2% arabinose, and grown for an additional 2 hours at 37°C. Cells were pelleted and lysed using the B-PER™ Bacterial Protein Extraction Reagent (Pierce) with added protease inhibitors (Roche, cOmplete™ ULTRA Tablets, Mini, EDTA-free; 1 tablets/10 mL), and lysozyme (Thermo Scientific) and DNaseI (Thermo Scientific) following the manufacturer’s instructions. Lysates were cleared by centrifugation and applied to a Strep-Tactin®XT 4Flow® column (IBA Lifesciences GmbH). The column was washed 5x with Buffer W (IBA Lifesciences GmbH), 1x with 1M NaCl, then 4x with Buffer W, followed by protein elution with Buffer BXT (IBA Lifesciences GmbH). Eluted proteins were separated on a 10% SDS-PAGE gel and a single band of ∼36 kDa was observed upon Coomassie blue staining. Protein concentration was determined using Qubit® Protein Assay Kit following manufacturer’s instructions.

Double-stranded DNA from the *yeiE* regulatory region (P*_yeiE_*), −240 to +59 relative to start codon (Fig S2), and from the *flhD* regulatory region (P*_flhD_*), −240 to +60 relative to start codon, was generated by colony PCR using Q5 DNA polymerase with primers and reaction conditions available in Table S2. Product size and purity were determined by agarose gel electrophoresis. DNA was purified using the PCR Purification Kit.

Electrophoretic gel mobility assays were performed with the Electrophoretic Gel Mobility Assay Kit (ThermoFisher), following the manufacturer’s instructions. Briefly, P*_yeiE_* DNA (40 ng) was mixed with the indicated concentrations of rYeiE in binding buffer (150 mM KCl, 0.1 mM dithiothreitol, 0.1 mM EDTA, 10 mM Tris, pH 7.4) for 20 minutes at room temperature. P*_flhD_* (40 ng) was similarly mixed with 800 nM rYeiE. Protein/DNA mixtures were separated on a 6% native polyacrylamide gel. Gels were stained with SYBR green and imaged using the ChemiDoc MP (BioRad) SYBR green gel setting. The experiment was performed on two occasions.

### YeiE quantification

The YeiE-3xFLAG chromosomal epitope tag was created using lambda-red recombinase with modification from the procedure described by Uzzau et al [51]. A 1635 bp G-block (Integrated DNA Technologies) was created based on the sequence of pSUB11 to incorporate the 3xFLAG sequence immediately 5’ to the *yeiE* stop codon, followed by a *frt*-flanked kanamycin-resistance cassette (Table S2) [51]. The G-block was transformed into 14028s containing pKD46 expressing lambda-red recombinase, as described [46]. Transformants were verified by PCR using primers flanking *yeiE*, moved into a clean genetic background by bacteriophage P22-mediated transduction, and the kanamycin-resistance cassette removed by flp-recombinase [44, 46].

Overnight cultures of bacteria encoding the *yeiE-3xFLAG* construct were diluted 1:100 in Lennox broth for 1.5 hours. Cultures were supplemented with 1 mM sulfate or sulfite or left untreated, and samples were collected at the time of treatment (t=0) and every 15 minutes thereafter for 1.5 hours. Cultures were pelleted, resuspended in PBS with protease inhibitors (Roche, cOmplete™ ULTRA Tablets, Mini, EDTA-free; 1 tablets/10 mL), normalized by OD₆₀₀, and lysed in 10X SDS-PAGE Loading Buffer at 100°C for 10 minutes. Equal volumes of each sample were loaded onto two 10% SDS-PAGE gels and proteins were separated at 150 V for 1 hour. For total protein quantification, one gel was stained with Coomassie blue and destained using a methanol–acetic acid solution. The second gel was transferred to a PVDF membrane (Turbo Transfer System, Bio-Rad) and western blot performed using mouse anti-FLAG IgG (1:1000 dilution, Life Technologies, Lot ZD393479A) as primary antibody and horse radish peroxidase conjugated goat anti-mouse IgG (1:10,000 dilution; Pierce, Lot 46-170-060115) as secondary antibody. Blots were developed with Clarity Max Western ECL Substrate following the manufacturer’s instructions (Bio-Rad).

Coomassie blue-stained gels were imaged using the Coomassie blue setting, and blots were imaged using colorimetric and chemiluminescent detection using automatic optimal settings (Bio-Rad Chemidoc MP). ImageLab 6.1 (Bio-Rad) volume tool was used to estimate total protein and YeiE-3xFLAG band intensity. To determine total protein, the total volume of all bands on Coomassie blue-stained gels were measured and compared with the sample of lowest mean intensity within a gel to generate a normalization factor for each sample. To determine YeiE-3xFLAG expression, the intensity of the ∼36kDa YeiE-3xFLAG band was multiplied by the total protein normalization factor for the sample to generate relative YeiE-3xFLAG. Relative YeiE-3xFLAG was normalized to time zero to calculate fold change in YeiE-3xFLAG expression. The experiment was repeated on three independent occasions.

### Mouse infection

Mouse infections were approved by the University of Wisconsin-Madison Institutional Animal Care and Use Committee (Protocol #V006255). The extraintestinal infection model was used as previously described with 8-10 week old female and male BALB/cJ mice obtained from Jackson Laboratories (strain 000651) [52]. Overnight bacterial cultures were washed twice in PBS, and mice were infected with approximately 10^4^ CFU of an equal mixture of the two competing strains by intraperitoneal injection. Mice were euthanized 48 hours postinfection. Organs were harvested, homogenized in PBS, serially diluted, and plated for enumeration. The competitive index (CI) between the WT and *ΔyeiE* mutant was determined as the ratio of WT to mutant bacteria after infection normalized to the inoculum ratio.

### Statistical analyses and data availability

For murine infection, log-transformed CI was compared between sexes by unpaired t-test with Holm-Šídák’s correction for multiple comparisons. Significant difference in sex-grouped CI was determined by Mann-Whitney test with Holm-Šídák’s correction for multiple comparisons. For growth curves, the area under the curve (AUC) was calculated from blanked OD_600_ data from each biological replicate. For single time point growth, the growth fold change was calculated as end CFU/mL normalized to starting CFU/mL. Strain and treatment differences in AUC and growth fold change were determined by one or two-way ANOVA with corrections for multiple comparisons. For comparisons of data over time, a repeated measures two-way ANOVA was used with corrections for multiple comparisons. For all analyses, P<0.05 was considered statistically significant. Analyses were performed using GraphPad Prism version 10.4.1.

## Supporting Information

**Fig S1.**
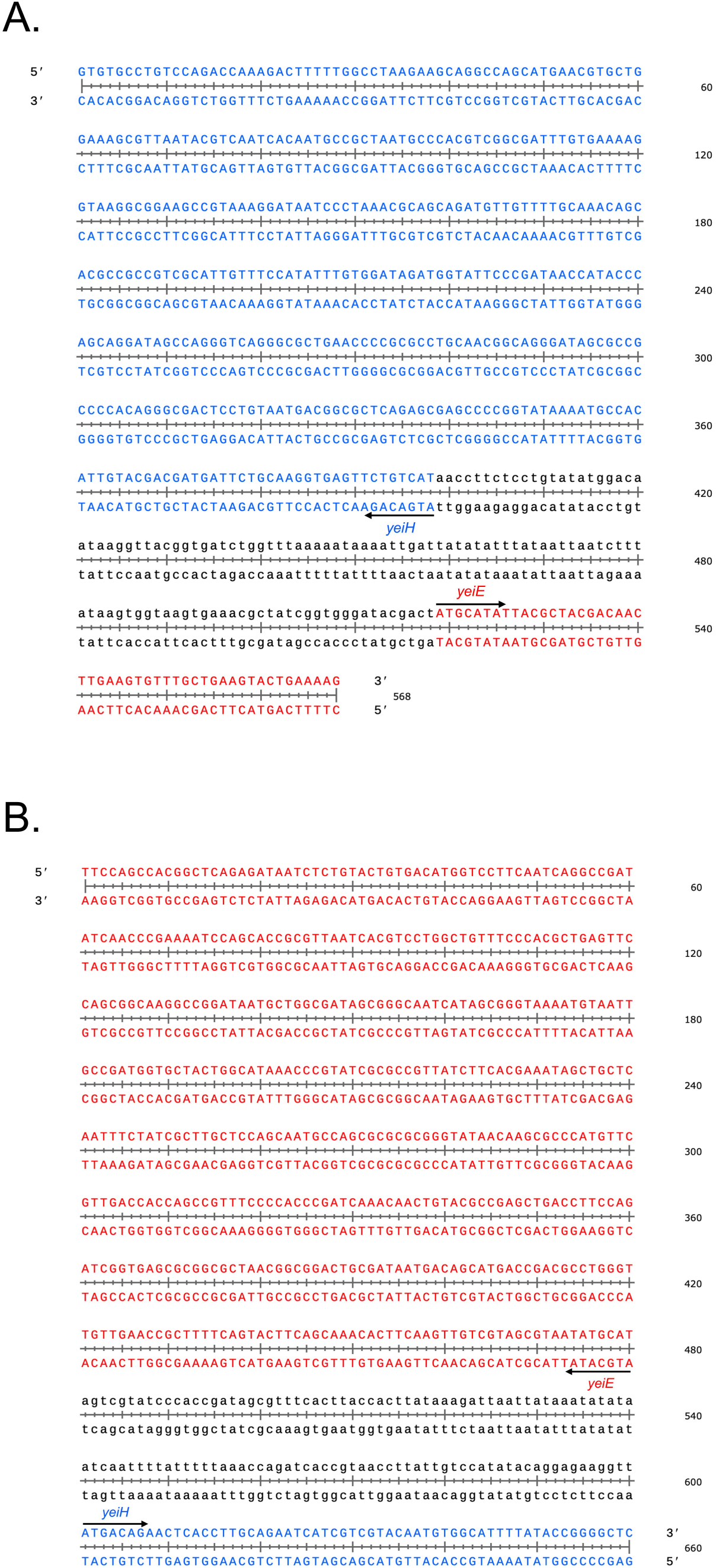
P*_yeiE_* and P*_yeiH_* sequences for pCS26-Pac.

**Fig S2.**
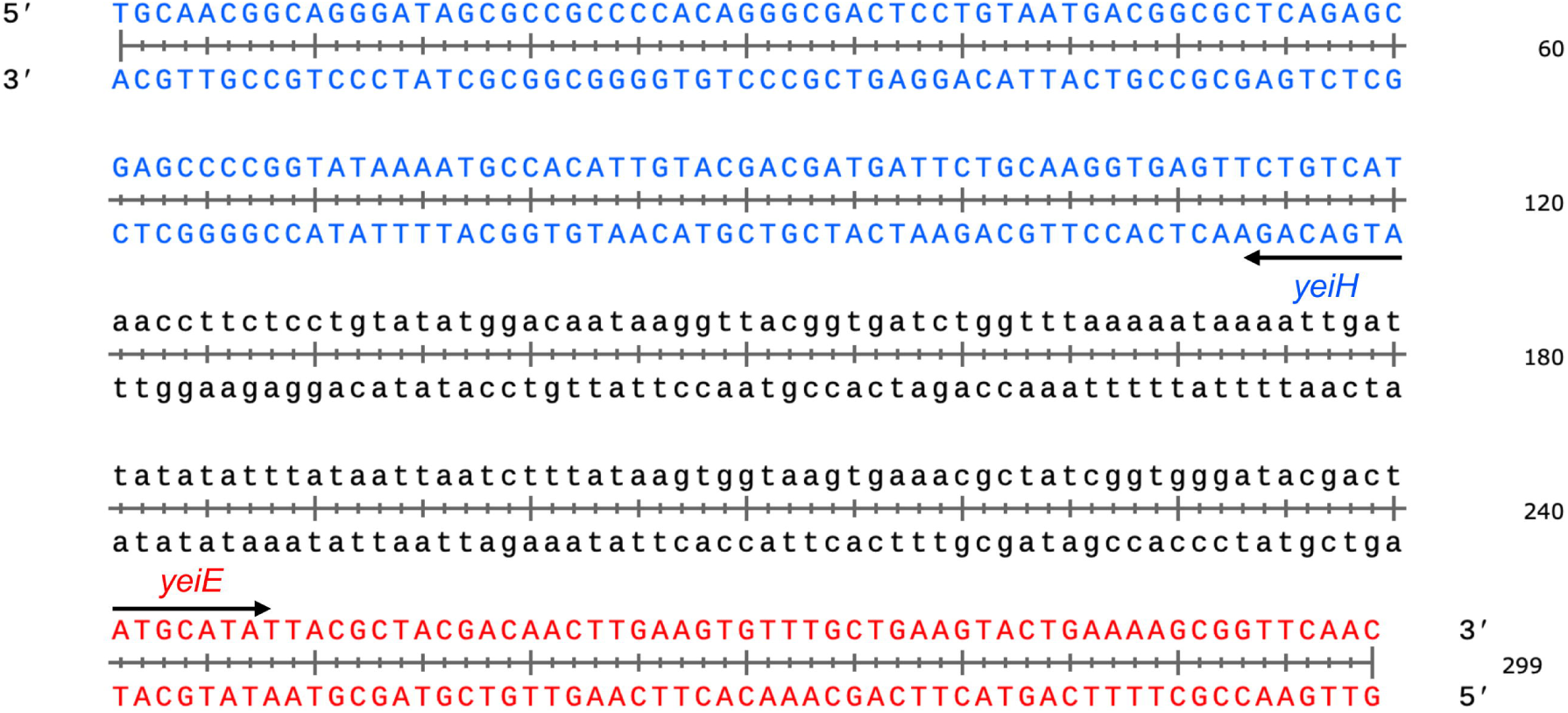
P*_yeiE_* DNA sequence for EMSA.

## Acknowledgements

This work was supported by the National Institute of Allergy and Infectious Diseases of the National Institutes of Health grant R21AI178071 to JRE and MM. ES was supported, in part, by the Food Research Institute Undergraduate Research Program in Food Safety. We thank Steffen Porwollik and Weiping Chu for supplying strains and Steffen for helpful comments on the manuscript.

