## Supplementary material for "YeiE regulates YeiH to implement sulfite stress resistance in *Salmonella enterica* serotype Typhimurium": Table S1

**Table S3: Raw data related to Fig 1.**

Experiment 1 - Females
WT vs ΔyeiE::Kan

| | Total count (Plate 1) | Total CFU/mL | Mutant count (Plate 2) | Mutant CFU/mL | WT CFU/mL | Inoculum dose (CFU/100ul) | Input Ratio (WT/mutant) |
|---|---|---|---|---|---|---|---|
| **Inoculum** | Nal | 1.62E+05 | Nal/Kan | 8.70E+04 | 7.50E+04 | 1.62E+04 | 0.862 |

| Organ | Mouse ID | Total count (Plate 1) | Total CFU/mL | Mutant count (Plate 2) | Mutant CFU/mL | WT CFU/mL (Total - Mutant) | Organ Output Ratio (WT/Mutant) | Output Ratio/Input Ratio | Log Output Ratio/Input Ratio |
|---|---|---|---|---|---|---|---|---|---|
| Spleen | M1 | Nal | 1.87E+04 | Nal/Kan | 4.30E+03 | 1.44E+04 | 3.35 | 3.88 | 0.59 |
| Spleen | M2 | Nal | 9.90E+03 | Nal/Kan | 2.04E+03 | 7.86E+03 | 3.85 | 4.47 | 0.65 |
| Spleen | M3 | Nal | 1.19E+04 | Nal/Kan | 3.40E+03 | 8.50E+03 | 2.50 | 2.90 | 0.46 |
| Spleen | M4 | Nal | 6.40E+02 | Nal/Kan | 1.10E+02 | 5.30E+02 | 4.82 | 5.59 | 0.75 |
| Spleen | M5 | Nal | 2.20E+04 | Nal/Kan | 6.20E+03 | 1.58E+04 | 2.55 | 2.96 | 0.47 |
| Liver | M1 | Nal | 2.07E+04 | Nal/Kan | 4.70E+03 | 1.60E+04 | 3.40 | 3.95 | 0.60 |
| Liver | M2 | Nal | 2.13E+04 | Nal/Kan | 3.20E+03 | 1.81E+04 | 5.66 | 6.56 | 0.82 |
| Liver | M3 | Nal | 1.18E+04 | Nal/Kan | 5.00E+03 | 6.80E+03 | 1.36 | 1.58 | 0.20 |
| Liver | M4 | Nal | 9.60E+02 | Nal/Kan | 1.60E+02 | 8.00E+02 | 5.00 | 5.80 | 0.76 |
| Liver | M5 | Nal | 2.20E+04 | Nal/Kan | 9.70E+03 | 1.23E+04 | 1.27 | 1.47 | 0.17 |

Experiment 2 - Males
WT vs ΔyeiE::Kan

| | Total count (Plate 1) | Total CFU/mL | Mutant count (Plate 2) | Mutant CFU/mL | WT CFU/mL | Inoculum dose (CFU/100ul) | Input Ratio (WT/mutant) |
|---|---|---|---|---|---|---|---|
| **Inoculum** | Nal | 2.56E+05 | Nal/Kan | 1.35E+05 | 1.21E+05 | 2.56E+04 | 0.894 |

| Organ | Mouse ID | Total count (Plate 1) | Total CFU/mL | Mutant count (Plate 2) | Mutant CFU/mL | WT CFU/mL (Total - Mutant) | Organ Output Ratio (WT/Mutant) | Output Ratio/Input Ratio | Log Output Ratio/Input Ratio |
|---|---|---|---|---|---|---|---|---|---|
| Spleen | M1 | Nal | 1.16E+06 | Nal/Kan | 1.71E+05 | 9.89E+05 | 5.78 | 6.47 | 0.81 |
| Spleen | M2 | Nal | 1.54E+04 | Nal/Kan | 3.00E+03 | 1.24E+04 | 4.13 | 4.62 | 0.66 |
| Spleen | M3 | Nal | 8.50E+05 | Nal/Kan | 1.39E+05 | 7.11E+05 | 5.12 | 5.72 | 0.76 |
| Spleen | M4 | Nal | 1.19E+05 | Nal/Kan | 1.59E+04 | 1.03E+05 | 6.48 | 7.25 | 0.86 |
| Spleen | M5 | Nal | 2.70E+05 | Nal/Kan | 5.60E+04 | 2.14E+05 | 3.82 | 4.27 | 0.63 |
| Liver | M1 | Nal | 3.10E+06 | Nal/Kan | 1.88E+05 | 2.91E+06 | 15.49 | 17.32 | 1.24 |
| Liver | M2 | Nal | 2.37E+04 | Nal/Kan | 5.90E+03 | 1.78E+04 | 3.01 | 3.36 | 0.53 |
| Liver | M3 | Nal | 1.04E+06 | Nal/Kan | 1.87E+05 | 8.53E+05 | 4.56 | 5.10 | 0.71 |
| Liver | M4 | Nal | 1.79E+05 | Nal/Kan | 2.90E+04 | 1.50E+05 | 5.17 | 5.78 | 0.76 |
| Liver | M5 | Nal | 7.00E+05 | Nal/Kan | 6.30E+04 | 6.37E+05 | 10.11 | 11.30 | 1.05 |
