## Supplementary material for "YeiE regulates YeiH to implement sulfite stress resistance in *Salmonella enterica* serotype Typhimurium": Table S2

**Table S2: Bacterial strains and plasmids**

**Bacteria**

| **Strain #** | **Genotype** | **Reference** |
|---|---|---|
| JE447 | WT (ATCC 14028s) | ATCC |
| JE2637 | ΔyeiE::frt | this study |
| JE3536 | JE2637+pWSK29 (Amp-R) | this study |
| JE3538 | JE2637+pWSK29::yeiE (Amp-R) | this study |
| JE1353 | ΔyeiE::kan (Kan-R) | Griewisch 2020 |
| JE3068 | ΔyeiH::kan (Kan-R) | this study |
| JE3297 | JE3068+pWSK29 (Kan-R, Amp-R) | this study |
| JE3299 | JE3068+pWSK29::yeiH (Kan-R, Amp-R) | this study |
| JE2616 | WT+pCS26-Pac (Kan-R) | this study |
| JE2629 | WT+pCS26-Pac-PyeiE::luxCDABE (Kan-R) | this study |
| JE2663 | JE2637+pCS26-Pac (Kan-R) | this study |
| JE2667 | JE2637+pCS26-Pac-PyeiE::luxCDABE (Kan-R) | this study |
| JE3501 | yeiE::3xFLAG | this study |
| JE3399 | ΔyeiH::frt | this study |
| HA420 | ATCC 14028s (spontaneous Nal-R) | Bogomolnaya 2008 |
| JE973 | HA420 ΔyeiE::kan (Nal-R, Kan-R) | Westerman 2021 |
| JE3193 | JE447+pCS26-Pac-PyeiH::luxCDABE (Kan-R) | this study |
| JE3199 | JE2637+pCS26-Pac-PyeiH::luxCDABE (Kan-R) | this study |
| JE3088 | 14028 ΔcysJ::kan | this study |
| JE2987 | 14028 ΔasrA::kan | this study |
| JE3148 | 14028 ΔcysJ::kanΔasrA::Cm (Kan-R, Cat-R) | This study |
| JE3466 | JE3399 ΔcysJ:kanΔasrA::Cm (Kan-R, Cat-R) | This study |
| JE3568 | JE2637 ΔcysJ::kanΔasrA::Cm (Kan-R, Cat-R) | This study |

**Plasmids**

| **Name** | **Resistance** | **Reference** |
|---|---|---|
| pCS26-Pac | Kan-R | Bjarnason 2003 |
| pBAD18Cm | Cat-R | Guzman 1995 |
| pWSK29 | Amp-R | Wang 1991 |
| pKD46 | Amp-R | Datsenko 1990 |
| pCP20 | Amp-R | Datsenko 1990 |
| pWSK29::yeiE | Amp-R | Westerman 2021 |
| pCS26-Pac::PyeiE-luxCDABE | Kan-R | This study |
| pCS26-Pac::PyeiH-luxCDABE | Kan-R | This study |
| pWSK29::yeiH | Amp-R | This study |
| pBAD18Cm::yeiE-strII | Cat-R | This study |
