## Supplementary material for "YeiE regulates YeiH to implement sulfite stress resistance in *Salmonella enterica* serotype Typhimurium": Table S3

**Table S3: Primers and synthetic nucleic acids**

**Cloning Primers**

| Forward | Reverse | Ta (˚C) | Extension time (sec) | Product size (bp) | Use |
|---|---|---|---|---|---|
| 5yeiE-strKpnIpBAD18<br>5' GTAACGGTACCATGCATATTACGCTACGACAACTTGAAG 3' | 3yeiE-strHindIIIpBAD18<br>5' GTAACAAGCTTTTATTTTTCGAACTGCGGGTGGCTCCAAGCGCTTTCACAGTAACTCAAAAAACGCTGC 3' | 65 | 60 | 916 | pBAD18-Cm-yeiE-strII |
| PyeiEFWD2XhoI<br>5' GTCCTCGAGGTGTGCCTGTCCAGACCAAAGAC 3' | PyeiEREV2BamHI<br>5' GTCGGATCCCTTTTCAGTACTTCAGCAAACAC 3' | 55 | 20 | 568 | pCS26-Pac-PyeiE-luxCDABE |
| PyeiHXhoIpCS26FWD<br>5' GGACTCGAGTTCCAGCCACGGCTCAGAG 3' | PyeiHBamHIpCS26REV<br>5' GTCGGATCCGAGCCCCGGTATAAAATGCCAC 3' | 66 | 35 | 660 | pCS26-Pac-PyeiH-luxCDABE |
| yeiH BamHI FWD<br>5' GATGGATCCCTATCGCTTGCTCCAGCAATGCCAG 3' | yeiH HindIII REV<br>5' GATAAGCTTCGCTATGATAACGGGTTAACAGGCGTG 3' | 62 | 30 | 1444 | pWSK29::yeiH |

**Primers for EMSA DNA fragments**

| Forward | Reverse | Ta (˚C) | Extension time (sec) | Product size (bp) |
|---|---|---|---|---|
| PflhDFWD2EMSA<br>5' CGTTGTATGTCACGAAGCTG 3' | PflhDREV EMSA<br>5' ATATGACAAATTGATGTCATAAATGTGTTTTAG 3' | 55 | 20 | 300 |
| PyeiEFWD2EMSA<br>5' TGCAACGGCAGGGATAGC 3' | PyeiEREV EMSA<br>5' GTTGAACCGCTTTTCAGTACTTC 3' | 55 | 20 | 299 |

**G-block sequence for *yeiE-3XFLAG***

5'
GAAACATCTTTCTAACGCGTTGCAGCGTTTTTTGAGTTACTGTGAAGACTACAAAGACCATGACGGTGATTATAAAGATCATGATATCGACTACAAAGATGACGACGATAAAT
AGTAATGTAGGCTGGAGCTGCTTCGAAGTTCCTATACTTTCTAGAGAATAGGAACTTCGGAATAGGAACTTCAAGATCCCCCACGCTGCCGCAAGCACTCAGGGCGCAAGG
GCTGCTAAAGGAAGCGGAACACGTAGAAAGCCAGTCCGCAGAAACGGTGCTGACCCCGGATGAATGTCAGCTACTGGGCTATCTGGACAAGGGAAATCGCAAGCGCAAA
GAGAAAGCAGGTAGCTTGCAGTGGGCTTACATGGCGATAGCTAGACTGGGCGGTTTTATGGACAGCAAGCGAACCGGAATTGCCAGCTGGGGCGCCCTCTGGTAAGGTTG
GGAAGCCCTGCAAAGTAAACTGGATGGCTTTCTTGCCGCCAAGGATCTGATGGCGCAGGGGATCAAGATCTGATCAAGAGACAGGATGAGGATCGTTTCGCATGATTGAAC
AAGATGGATTGCACGCAGGTTCTCCGGCCGCTTGGGTGGAGAGGCTATTCGGCTATGACTGGGCACAACAGACAATCGGCTGCTCTGATGCCGCCGTGTTCCGGCTGTCA
GCGCAGGGGCGCCCGGTTCTTTTTGTCAAGACCGACCTGTCCGGTGCCCTGAATGAACTGCAGGACGAGGCAGCGCGGCTATCGTGGCTGGCCACGACGGGCGTTCCTTG
CGCAGCTGTGCTCGACGTTGTCACTGAAGCGGGAAGGGACTGGCTGCTATTGGGCGAAGTGCCGGGGCAGGATCTCCTGTCATCTCACCTTGCTCCTGCCGAGAAAGTAT
CCATCATGGCTGATGCAATGCGGCGGCTGCATACGCTTGATCCGGCTACCTGCCCATTCGACCACCAAGCGAAACATCGCATCGAGCGAGCACGTACTCGGATGGAAGCC
GGTCTTGTCGATCAGGATGATCTGGACGAAGAGCATCAGGGGCTCGCGCCAGCCGAACTGTTCGCCAGGCTCAAGGCGCGCATGCCCGACGGCGAGGATCTCGTCGTGA
CCCATGGCGATGCCTGCTTGCCGAATATCATGGTGGAAAATGGCCGCTTTTCTGGATTCATCGACTGTGGCCGGCTGGGTGTGGCGGACCGCTATCAGGACATAGCGTTGG
CTACCCGTGATATTGCTGAAGAGCTTGGCGGCGAATGGGCTGACCGCTTCCTCGTGCTCTACGGTATCGCCGCTCCCGATTCGCAGCGCATCGCCTTCTATCGCCTTCTTGA
CGAGTTCTTCTGAGCGGGACTCTGGGGTTCGAAATGACCGACCAAGCGACGCCCAACCTGCCATCACGAGATTTCGATTCCACCGCCGCCTTCTATGAAAGGTTGGGCTTC
GGAATCGTTTTCCGGGACGCCGGCTGGATGATCCTCCAGCGCGGGGATCTCATGCTGGAGTTCTTCGCCCACCCCAGCTTCAAAAGCGCTCTGAAGTTCCTATACTTTCTAG
AGAATAGGAACTTCGGAATAGGAACTAAGGAGGATATTCATATGAGGTGAAGAGGAGGGCTGCCCGGGCGGCATCGCCATTTTTAG 3'
